# Peptides that Mimic RS repeats modulate phase separation of SRSF1, revealing a reliance on combined stacking and electrostatic interactions

**DOI:** 10.1101/2022.10.24.511151

**Authors:** Talia Fargason, Naiduwadura Ivon Upekala De Silva, Erin King, Zihan Zhang, Trenton Paul, Jamal Shariq, Steve Zaharias, Jun Zhang

## Abstract

Phase separation plays crucial roles in both sustaining cellular function and perpetuating disease states. Despite extensive studies, our understanding of this process is hindered by low solubility of phase-separating proteins. One example of this is found in SR proteins. These proteins are characterized by domains rich in arginine and serine (RS domains), which are essential to alternative splicing, *in vivo* phase separation, and a low solubility that has made these proteins difficult to study for decades. Here, we solubilize the founding member of the SR family, SRSF1, by introducing a peptide mimicking RS repeats as a co-solute. We find that this RS-mimic peptide forms interactions similar to those of the protein’s RS domain. Both interact with a combination of surface-exposed aromatic residues and acidic residues on SRSF1’s RNA Recognition Motifs (RRMs) through electrostatic and cation-pi interactions. Analysis of RRM domains spanning the human proteome indicates that RRM domains involved in phase separation have more exposed aromatic residues and that in phase-separating proteins containing RS repeats, such residues are frequently surrounded by acidic residues. In addition to opening an avenue to previously unavailable proteins, our work provides insight into how SR proteins phase separate and participate in nuclear speckles.

## INTRODUCTION

Liquid-liquid phase separation underpins the formation of membraneless organelles, such as nucleoli(1), P-bodies(2), stress granules(3), cajal bodies(4), and nuclear speckles(5). The integrity of such organelles is maintained by interactions between biomolecules that form condensates, or liquid droplet-like structures, in which the local concentration of individual components is higher than the surrounding environment (6). These condensates cluster relevant molecules together to facilitate interactions while allowing rapid material exchange(6-9). Mounting evidence has revealed roles of phase separation in modulating reaction kinetics, enzyme catalysis, and binding specificity(10-14).

The protein SRSF1 (Serine/Arginine-Rich Splicing Factor 1, also known as ASF/SF2) is essential for the early-stage assembly of the spliceosome (15,16). Several *in vivo* studies have shown that SRSF1 is found in condensates (5,17-22). Aberrant condensation behaviors have also been observed in disease states (20-22). SRSF1 belongs to the Ser/Arg-rich protein family (SR proteins), which contains 12 members possessing one to two structured RNA-recognition motifs (RRMs) and a repetitive Arg/Ser repeat region (RS domain)(23-25). The RS region is also found in the much larger family of SR-like proteins, which contain the repetitive RS regions but not the other structural features (26,27). Two SR-like proteins, SRRM2 (serine-arginine rich repetitive matrix protein 2) and SON, are essential for the formation and structural maintenance of the membraneless organelles nuclear speckles (28,29). Like many other splicing factors, SRSF1 modulates trafficking to the speckles(30). It has been demonstrated that SRSF1 is found in nuclear speckles when its RS domain is partially phosphorylated but that hyperphosphorylation causes the protein to leave nuclear speckles(31,32). It is therefore evident that RS domains play an important role in the organization of nuclear speckles. However, an understanding of the nature of that interaction has been evasive due to a difficulty solubilizing the proteins involved. As with all 12 SR proteins and many speckle components, obtaining soluble SRSF1 has been an imposing challenge for decades, and this has substantially hindered our understanding of the functions of these proteins and of nuclear speckles as a whole(23).

Phase separation is frequently mediated by repetitive sequences. Here, we find this to be the case for the protein SRSF1, whose phase separation is dependent on its RS repeats. Our bioinformatic analysis reveals a correlation of RS repeats with a tendency to phase separate. We successfully solubilize SRSF1 using short peptides that mimic RS repeats. Our success in solubilizing SRSF1 provides an unprecedented opportunity to elucidate the mechanism by which RS repeats interact with SRSF1. We find that this increase in solubility is due to a competition between the peptide and RS domains for the same binding sites on RRM domains. We further discover that acidic residues and aromatic residues from SRSF1 RRMs interact with the RS region through salt bridges and cation-pi interactions. We find that many of the RRM sites interacting with RS repeats are conserved among the SR protein family. Further, we find that RRM domains belonging to phase-separating proteins have more solvent-accessible aromatic residues than RRM domains of proteins that have not been found in condensates. In proteins containing RS repeats and proteins that localize to nuclear speckles, these solvent exposed aromatic residues are more likely to be surrounded by neighboring acidic residues. These findings provide insight into how interactions between SR and SR-like proteins may occur within membraneless organelles. They also allow us to predict how the nature of these interactions might change when the RS domain becomes phosphorylated.

## RESULTS

### RS repeats are abundant in the human proteome and associated with phase separation

RS repeats are often found in SR proteins and SR-like proteins. The serine residues in RS repeats are frequently phosphorylated, a process which regulates the functions of RS repeats. To quantify the abundance of RS repeats in the human proteome, we systematically searched for uninterrupted repeats that were 2 to 8 amino acids in length (Fig. 1A) and tested whether the length of RS repeats was correlated with protein condensates. Proteins were defined as being in condensates if they were listed in any of three available phase separation databases (PhaSepDB, LLPSDB, and DrLLPS) (33-35). Because both RRM domains and RS repeats have been associated with phase separation in previous studies, we separated proteins containing RRM domains in this analysis from those that did not(36,37). We found that the chance that a protein was found in condensates increased with the length of RS repeats it harbored regardless of whether the protein contained an RRM domain (Fig. 1A). Proteins possessing both RS repeats and RRM domains were more likely to be found in condensates (Fig. 1A, blue bars).

**Figure 1.**
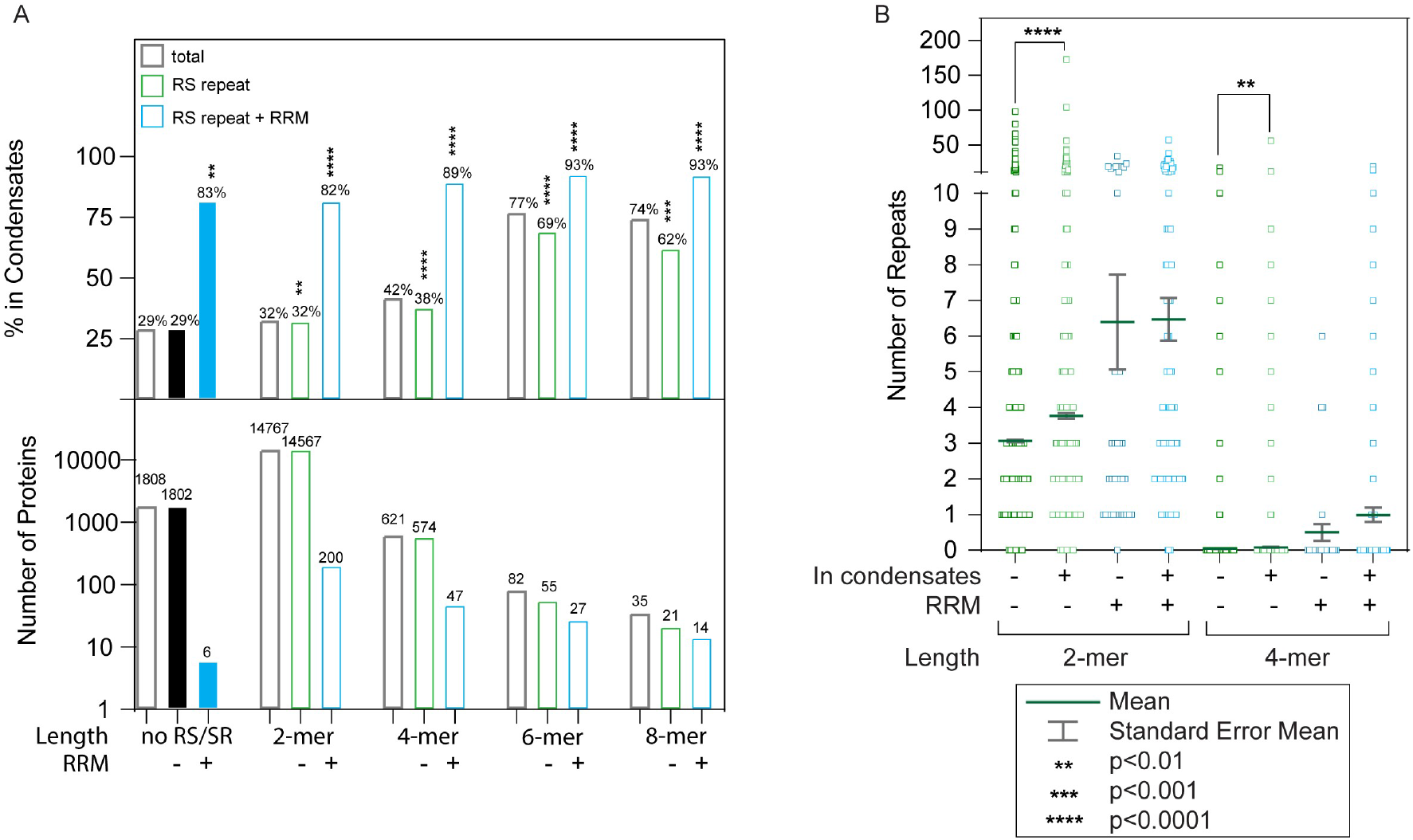
A combination of RS repeats and RRM domains is highly correlated with appearance in condensates. (A) Increased RS repeat length leads to an increased likelihood of appearance in condensates. Percentage of proteins possessing indicated properties that appear in one of three major phase separation databases. The *p*-values were obtained using Fisher’s test for exactness, comparing values to those found for proteins possessing neither RS repeats not RRM domains (black column). (B) Correlation between number of 2-mer RS and 4-mer RS repeats with appearance in condensates. Proteins found in condensates are more likely to have a greater number of RS dipeptide and tetra-peptide repeats. The *p*-values presented were obtained using the Mann-Whitney test.

Only 6 proteins with RRM domains lacked a single RS dipeptide. Although five of them have been found in condensates, it is difficult to draw conclusions about the effect of RRM domains on phase separation based on this. Proteins that contained both RRM domains and RS repeats are more likely to phase separate than those possessing neither component. Among proteins with an RRM and at least one 4-mer RS repeat, the likelihood of appearance in condensates was 89%, and this trend became more pronounced as the repeat length was increased (Fig. 1A).

Many proteins contain multiple copies of 2-mer RS or 4-mer RS repeats. One extreme example is nuclear speckle scaffolding protein SRRM2, which has 56 4-mer RS repeats, more than any other protein (Fig. S1). We found that the number of RS repeats a protein harbors also affects its likelihood to be found in condensates (Fig. 1B). Proteins in condensates have more copies of 2-mer or 4-mer RS repeats than those not in condensates on average (Fig. 1B).

### SRSF1 can be solubilized using peptides that mimic its RS region

The correlation of RS-repeats with phase separation is consistent with observations that many RS-containing proteins have low solubility *in vitro*. For example, up to this point, none of the full-length SR proteins have been obtained in concentrations suitable for biophysical/biochemical or structural characterization, although the founding member of the family, SRSF1, was identified more than three decades ago (38-40).

To overcome this obstacle in investigating SR and SR-like proteins, we aimed to develop a new purification and solubilization method using full-length SRSF1. The high Arg composition in these proteins inspired us to use high concentrations of Arg amino acid in our protocol to purify and solubilize SRSF1. We found that 0.8 to 1 M of Arg was able to solubilize all SRSF1 constructs during the purification procedure (details in the method section). However, the high ionic strength of Arg at this concentration range is unsuitable for many analytical methods, such as NMR and binding assays.

We predicted that we could solubilize phase-separating proteins using peptide co-solutes that mimic these repeats to compete with inter- and intra-molecular interactions (Fig. 2A). We tested this concept on SRSF1, which contains RS repeats of 16, 5, and 6 amino acids, respectively (Fig. 2B). Serine residues in these regions can be phosphorylated, resulting in an alternation of positive and negative charges (Fig. 2B). To solubilize SRSF1 in its unphosphorylated and phosphorylated forms, we therefore tested peptides of varying lengths to mimic unphosphorylated RS (RS), and phosphorylated RS (DR, and ER) repeats (Fig. 2B). Here, using the purified protein, we found that SRSF1 phase separated at concentrations lower than 300 nM in a phosphate buffer (Fig. 2C, 2D). This was also the case when the protein was diluted into 90 mM KCl (Fig. 2E).

**Figure 2:**
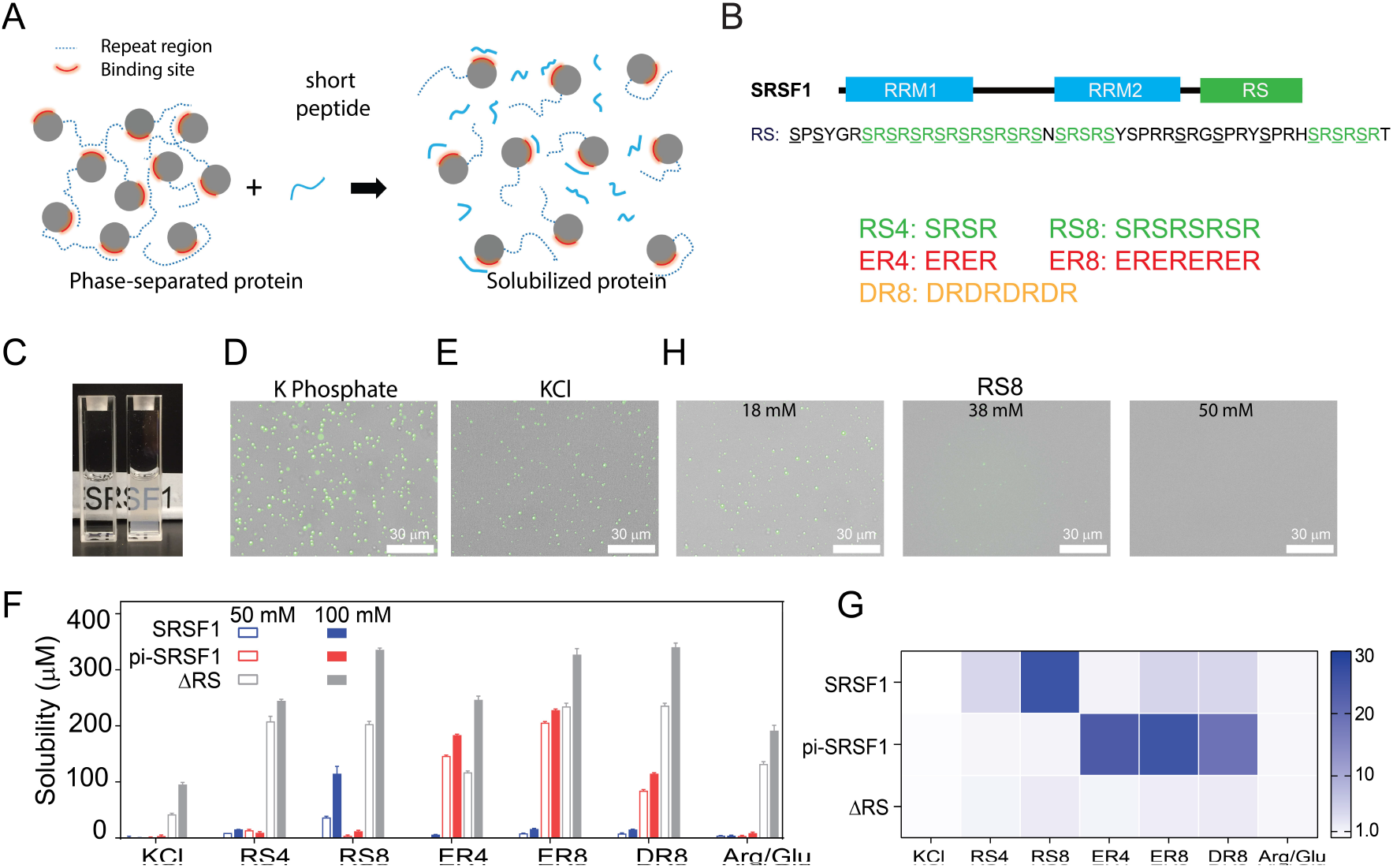
SRSF1 phase separation can be reduced using peptides that best mimic its RS repeats in their respective phosphorylation states. (A) Schematic illustration of solubilizing phase-separating proteins using short peptides. Short peptides compete with RS repetitive regions, disrupting phase separation. (B) Domain architecture of SRSF1. The underscored serine residues in the SRSF1 RS can be phosphorylated, and the phosphorylated RS can be mimicked by ER and DR repeats. Short peptide co-solutes used in this study are shown below. (C) Phase separation of SRSF1. The left cuvette is SRSF1 solubilized in the RS8 peptide, and the right cuvette is SRSF1 in 140 mM potassium phosphate, pH 7.4, 10 mM NaCl. The fluorescence image of 288 nM unphosphorylated SRSF1 in phosphate buffer (D), KCl buffer (E). SRSF1 is labeled with Alexa488 at N220C. (F) SRSF1 solubility using 50 mM or 100 mM of peptide as indicated. (G) Ratio of solubility in peptides to solubility in 100 mM Arg/Glu as determined in panel D. (H) The RS8 peptide can reduce phase-separation droplets of SRSF1.

To quantify protein solubility, we used ammonium sulfate precipitation followed by resuspension of the proteins in peptide-containing buffers (Fig. 2F). This approach to measuring protein solubility has been used in many studies (41,42). If mimicking the repetitive sequences with the peptides helps resolve phase separation, we expect to see a more dramatic solubility increase when the peptide co-solutes most closely mimic the repetitive sequences of the proteins. To this end, we measured solubility of unphosphorylated full-length SRSF1, hyper-phosphorylated SRSF1 (pi-SRSF1), and RS-truncated SRSF1 (ΔRS) (Fig. 2F). We found that whether phosphorylated or not, full-length SRSF1 was essentially insoluble in the 50 mM and 100 mM KCl control buffers (Fig. 2F). Previous studies have found that truncation of the RS domain increases protein solubility(24). This suggests that the RS domain is responsible for the protein’s low solubility. To verify this, we measured the solubility of ΔRS and found that it was overall more soluble than full-length SRSF1 in all tested buffers (Fig. 2F). Although an Arg/Glu mixture has been reported to promote protein solubility(43), Arg/Glu at 100 mM provided only a limited solubilizing effect for full-length SRSF1 (Fig. 2F). Among the peptides we tested, only RS8 dramatically increased solubility of unphosphorylated SRSF1 (from 0.6 ± 0.29 μM in 100 mM KCl to 120 ± 12 μM with 100 mM RS8). Consistent with our hypothesis, all tested peptides had a less dramatic solubilizing effect on ΔRS, likely because it does not contain the repeat sequences that the peptides are designed to mimic. Hyper-phosphorylation of an RS domain converts the region into a basic acidic repeat resembling ER and DR repeats. To mimic a phosphorylated RS domain, we tested the solubilizing effects of ER and DR peptides. ER8 increased the solubility of hyper-phosphorylated SRSF1 (pi-SRSF1) more than other peptides. DR8 and ER4 also had substantial, albeit less pronounced, solubilizing effects (Fig. 2F). The preference for ER8 may be due to the fact that glutamic acid resembles phosphoserine more than aspartic acid. This confirmed our hypothesis that the solubilizing effect was more notable with peptides most closely resembling the proteins’ own repeats.

These trends are clearer when the effect of the peptides on solubility is normalized by the solubility in 100 mM Arg/Glu (Fig. 2G). Whereas peptides designed to mimic the repetitive constructs produce as much as a 30-fold increase in solubility relative to Arg/Glu, the solubility increase of ΔRS only reaches about a 2-fold difference (Fig. 2G). In accordance with our solubility tests, we found that using increasing concentrations of the RS8 peptide reduced the number of liquid-like droplets in solution (Fig. 2H).

We also tested the solubilizing effect of these peptide co-solutes on two other RNA-binding proteins, Nob1 and Nop9. These peptides increased solubility of Nob1 and Nop9 by 50-60% and 2-10%, respectively (Fig. S2A). Nob1 contains unstructured regions rich in basic and acidic-basic residues (Fig. S2B) and has an increased solubility despite its unstructured regions having a lower homology to the tested peptides. Nop9 does not have such unstructured sequence regions (Fig. S2C). Consistent with our hypothesis, the tested peptides have a moderate solubilizing effect on Nob1 whereas they have limited or no effect on Nop9. These results for SRSF1 constructs, Nob1, and Nop9 suggest that repetitive peptides improve solubility for proteins that have similar sequences.

### Peptide co-solutes are compatible with NMR experiments and binding assays

Ionic co-solutes usually increase the dielectric constant of a sample and complicate NMR data acquisition, producing difficulty in probe tuning/matching, elongation of pulse width, and reduction of sensitivity(44,45). This imposes a considerable obstacle, as NMR is one of the few methods that provides an atomic level description of the dynamic interactions of phase-separating proteins. This adverse effect can be quantified by the elongation of the pulse width, which is inversely proportional to the signal sensitivity(44). For example, increasing KCl concentration from 100 mM to 400 mM elongates the pulse width by about 50% on the NMR probe used in this study (Fig. 4A). In contrast to the effect of KCl, peptide co-solutes did not significantly elongate the pulse width (Fig. 4A). Whereas the 800 mM Arginine buffer used to solubilize SRSF1 during purification increased the pulse width to 16.97 µs, a combination of peptide and arginine had a less pronounced effect on the pulse width (Fig. S4A). This is likely due to the low mobility of short peptides compared with salts or free amino acids(45).

We expect the peptides to compete with homotypic inter-molecular interactions (interactions between SRSF1 molecules) to solubilize the protein, but the competition should not be strong enough to abolish binding or disrupt protein structure. With the peptide co-solutes, we were able to obtain high quality NMR spectra for both unphosphorylated and phosphorylated SRSF1. The TROSY-HSQC overlay in Fig. 3B is consistent with the expected presence of both globular domains that show a higher level of dispersion and disordered regions with proton shifts in the 8.0-8.5 ppm range. Although we were able to solubilize unphosphorylated and phosphorylated SRSF1 in these respective buffers, a buffer that solubilizes both proteins was desired to allow direct comparison of the two proteins and facilitate NMR assignment. We found that a buffer of 100 mM ER4 mixed with 400 mM Arg/Glu, pH 6.4, maintained structure and solubilized both unphosphorylated and phosphorylated SRSF1 constructs at concentrations above 350 µM (Fig. S3). It also weakened but did not abolish binding of SRSF1 constructs to an RNA ligand (Fig. S4B) and resulted in a pulse width of 15.04 µs (Fig. S4A), significantly shorter than that observed for 800 mM Arg/Glu. Using this buffer, we assigned the backbone for unphosphorylated SRSF1(Fig. 4C, D). This accomplishment enabled us to investigate the mechanism by which repetitive peptides solubilize SRSF1. To this end, we selected RS8 and unphosphorylated SRSF1 for further study.

**Figure 3:**
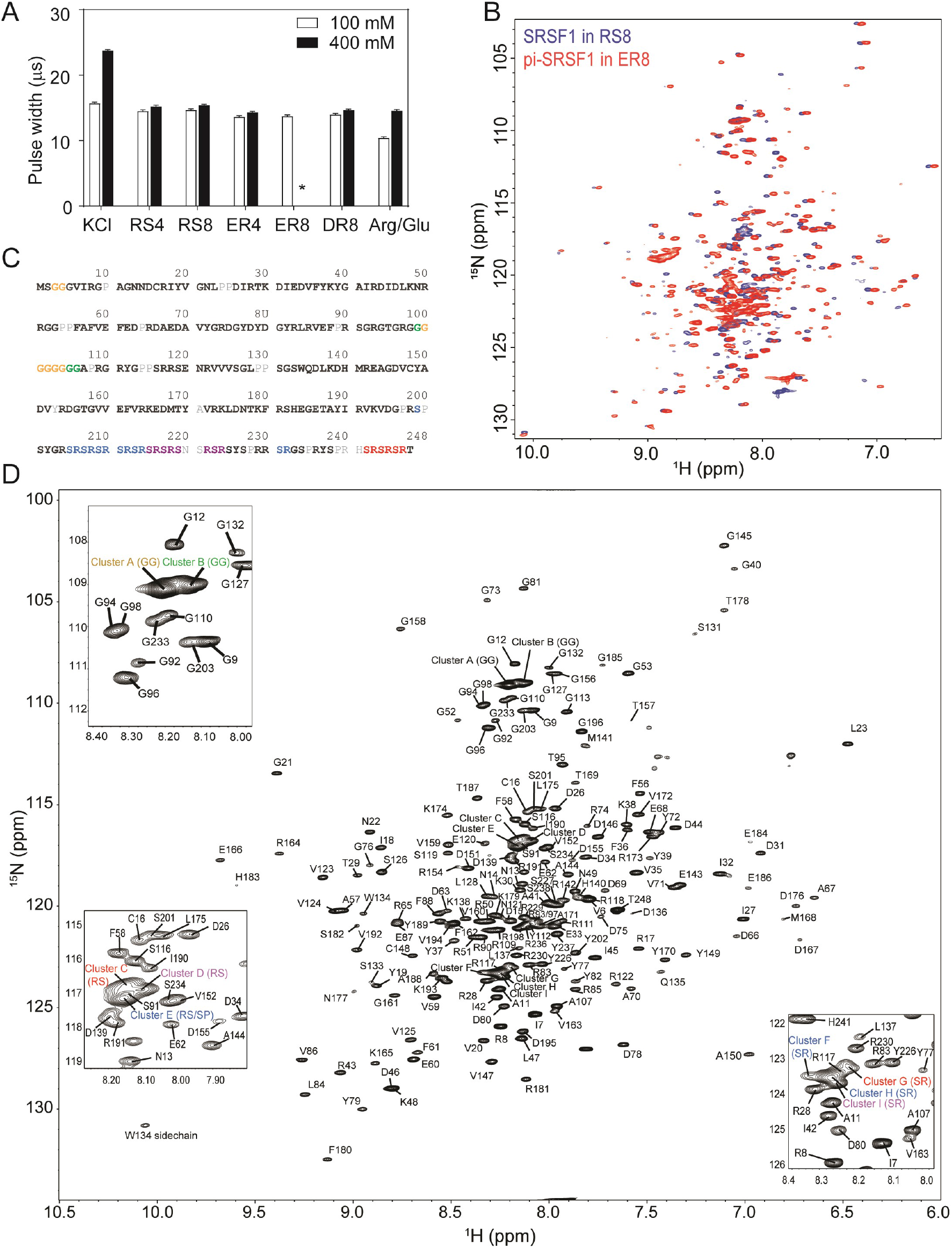
Short peptides are compatible with NMR experiments. (A) NMR 90-degree pulse width. ER8 is insoluble at 400 mM and therefore its pulse width could not be determined, as indicated by *. (B) ^15^N-TROSY-HSQC overlay of SRSF1 in 100 mM RS8 and phosphorylated SRSF1 (pi-SRSF1) in 100 mM ER8. (C) Assigned residues in the SRSF1 protein sequence. Black bold fonts indicate non-overlapping residues. Gray fonts indicate unassigned residues. Color fonts indicate amino acids assigned to clusters. (D) Assignment of the SRSF1 amide groups.

**Figure 4:**
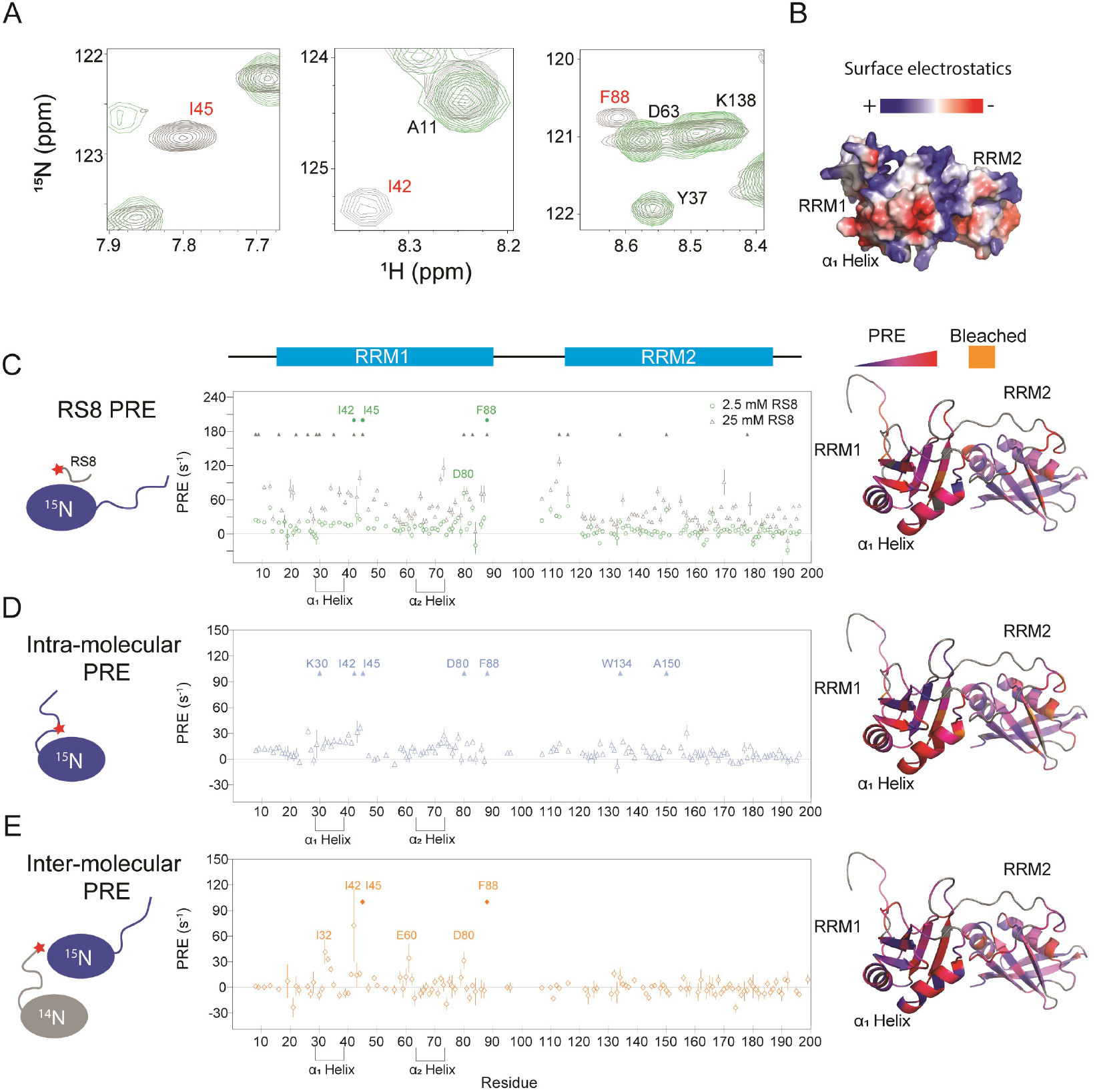
SRSF1 residues involved in interactions with the RS8 peptide are similar to those found in intra-, and homotypic inter-molecular interactions with the RS region. (A) ^15^N-TROSY-HSQC overlay of SRSF1 in 50 mM diamagnetic (gray) and 2.5 mM paramagnetic RS8 (green). The intensities of residues close to the probe become diminished. Bleached residues (indicated by red type) came in such close contact with RS8 that their intensities were diminished before the first observation timepoint (additional information in the methods section). The full spectra are shown in Fig. S5. (B) Electrostatic surface of SRSF1 RRM1 and RRM2. The α1 helix on RRM1 has a large negatively charged surface area, and RRM1 possesses overall more negative charge. (C) PRE values induced by 2.5 or 25 mM paramagnetic RS8. (D) Intra-molecular PRE produced by the MTSL-labeled RS region (N220C). (E) Inter-molecular PRE produced by the MTSL-labeled NMR-inactive SRSF1 (T248C). The filled symbols indicate bleached residues. Gray in the molecular graphics on the right indicates residues whose PRE values are unavailable due to peak overlap or an inability to assign them. PyMOL molecular graphics were prepared using Xplor-NIH (see methods section for more information).

### Acidic and exposed aromatic residues of SRSF1 RRMs are responsible for the interactions with RS repeats that lead to phase separation

Mimic peptides were able to provide us with control over the critical point for SRSF1 phase separation, enabling us to obtain a backbone assignment in the solution state(Fig. 3). According to our hypothesis, the mimic peptide should provide transient competition for contacts with the protein’s repetitive sequence. Inter- and intra-molecular interactions should still occur under these conditions. However, they should be weakened enough to prevent droplet formation, providing us with a stable sample that can be used to study the intermolecular interactions that lead to the initiation of phase separation. To verify that this was the case, we performed a series of paramagnetic relaxation enhancement NMR experiments to probe peptide, intramolecular, and homotypic intermolecular interactions.

To locate the RS-mimic peptide interacting sites, we labeled RS8 with a paramagnetic probe (MTSL) and mixed it with SRSF1 (Fig. 4A, 4C, and S5A). The paramagnetic group attenuates resonance intensities of residues by enhancing relaxation in a distance-dependent manner(46). PRE is suitable for probing transient interactions, including the weak interactions between co-solutes and macromolecules(46,47) and the intermolecular interactions that proceed phase separation(48,49). A higher PRE value indicates that peptides come closer to the residue analyzed. We found that paramagnetic RS8 interacted primarily with RRM1 residues (Fig. 4A, 4C, and Fig. S5). This is consistent with the fact that RRM1 (pI=4.7) is more acidic than RRM2 (pI=6.9), with regions of high negative charge on its two helices (Fig. 4B). Dramatically perturbed sites were clustered on electronegative and aromatic sites, with the sequence D^31^IED on the α1-Helix and D^44^ID on the neighboring β-sheet being particularly perturbed (Fig. 4C). Other hotspots included D^80^GYR and E^87^F on loops neighboring the α_1_ and α2 helices, respectively,D^66^AED on the α_2_ helix, and RRM2 residues W134 and A^150^DVYR (Fig. 4C).

According to our hypothesis, the RS8 peptide should provide transient competition with the RS domain without abolishing inter- and intra-molecular interactions (Fig. 2A). To locate the intra-molecular interacting sites, we separately introduced the MTSL label to the center of the RS domain (N220C, Fig. 4D, S5B) and the C-terminal end (T248C, Fig. S6B). We found that labeling at the center of the RS domain produced the most notable perturbations (Fig. 4D). To estimate the background PRE resulting from stochastic collisions, we also collected PRE data for SRSF1 mixed with the same concentration of MTSL alone as a control (Fig. S6A). Subtracting the background PRE does not significantly change the perturbation pattern (Fig. S6D,S6E). To verify that intermolecular interactions were not contributing to the measured intramolecular PRE, we performed a control PRE measurement by mixing an equal amount of MTSL-labeled HSQC-undetectable (^14^N) SRSF1 and ^15^N-SRSF1 with no cysteines (Fig. S6C). At this total concentration of 200 µM and with this labeling site (N220C-MTSL), we did not see a significant intermolecular contribution to the PRE signal. The relative strengths of the spectra are readily observed when the intermolecular interactions under these conditions are subtracted from the intramolecular interactions (Fig. S6F).

To locate the inter-molecular interacting sites, we placed MTSL on the C-terminal end of the protein and doubled the concentration to a total of 400 µM protein, maintaining a 1:1 ratio of HSQC-undetectable SRSF1 with MTSL and ^15^N-labeled SRSF1 possessing no cysteines (Fig. 4E, S5C). With this experimental design, only inter-molecular interactions resulted in PRE on the ^15^N-labeled SRSF1. Consistent with our hypothesis, the intra- and inter-molecular PRE patterns are similar to the perturbations from paramagnetic RS8.

To gain an atomic-level picture of the interactions between RS8 and SRSF1, we constructed models with the program Xplor-NIH using the PRE values as restraints(50,51). We further optimized these models using molecular dynamics simulations (Fig. 5, S7). Representative images are displayed in Fig. 5. The hotspot D^31^IED on the α1 helix was found to be able to provide electrostatic contacts for multiple interactions, including hydrogen bonding with bleached isoleucine residues I42 and I45 (Fig. 5A). In the region surrounding W134, an electrostatic interaction with D151 was found to enable a peptide arginine to orient parallel to the aromatic face of W134, forming cation-pi stacking interactions (Fig. 5B). Simultaneous cation-pi stacking interactions were also observed in the regions surrounding D80 and Y79 (Fig. 5C) as well as F88 and E68 (Fig. 5D).

**Figure 5.**
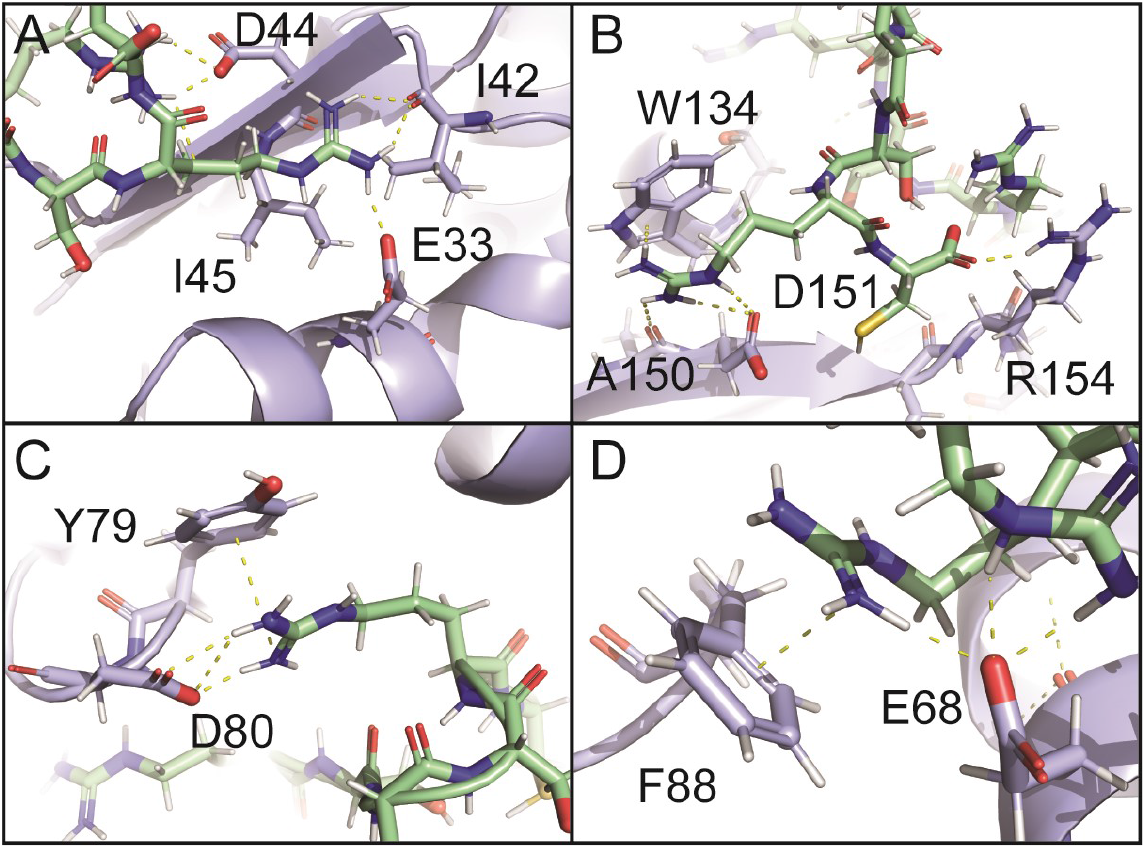
Electrostatic and cation-pi interactions are responsible for intermolecular interactions. Molecular dynamics simulation of SRSF1 with four RS8 peptides.

### RRM domains involved in phase separation possess more exposed aromatic residues

We were curious whether interactions found in SRSF1 were conserved across the SR protein family. To this end, we used the program ClustalX(52) to align the RRM1 sequences of SR proteins and found that interaction hotspots on α1 and α2 helices of RRM1 domains were conserved in most SR proteins (Fig. 6A). The electronegative charge was most highly conserved in the α2 helix and the neighboring loop, while in the α1 helix, acidic residues were replaced by arginine residues in some SR proteins. Interestingly, these regions are separate from the RNA-binding sites of the RRM domains, suggesting a role outside of RNA recognition (Fig. 6A). This analysis suggests that the phase-separating mechanism we revealed for SRSF1 could be applicable to many other members of the SR family.

**Figure 6.**
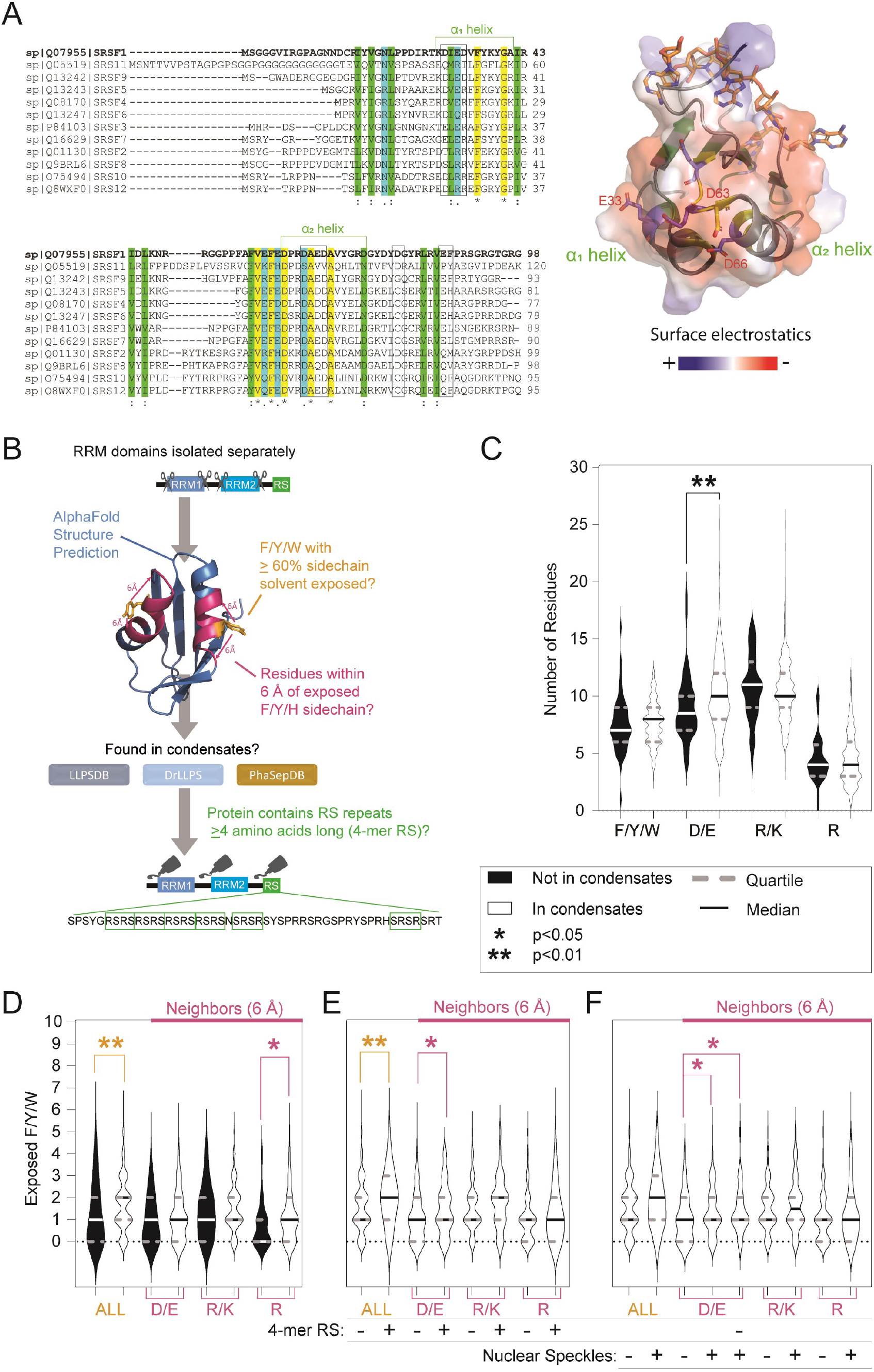
RRM domains found in condensates possess distinct features. (A) ClustalX alignment of RRM1 domains of the SR protein family, where yellow indicates identical amino acids, green and blue indicate conserved residues. Black boxes indicate PRE hotspots. Structure of RNA-bound SRSF1 RRM1 was obtained from PDB ID 6HPJ. Transparent electrostatic surface is displayed. Conserved electronegative residues opposite the RNA binding pocket are shown in sticks. (B) Schematic of the approach to the bioinformatics search. (C) Analysis of the amino acid composition of 365 RRM domains by residue and charge. Solid lines of violin plots represent medium values while dashed lines are upper and lower quartiles. (D) Analysis of solvent accessible aromatic residues. Solvent accessible aromatic residues were identified using the program Xplor-NIH and defined as those with ≥ 60% of sidechain area accessible to solvent. Basic (R/K), acidic (D/E), and R residues within 6 Å of solvent accessible aromatic residues were counted for RRMs in and not in condensates. (E) Examination of the RRM domains of proteins in condensates based on whether the proteins do or do not contain 4-mer RS repeats. “-” and “+” in the second row below the plot indicates 4-mer RS repeats are absent or present in these proteins. (F) Comparison of the RRM domains of proteins found in nuclear speckles vs other condensate types. “-“ and “+” in the third row below the plot indicates proteins have not or have been found in nuclear speckles. Column 5 examines the RRM domains of proteins that are found in nuclear speckles but do not contain their own RS4 repeats. RRM domains were classified as being in condensates if they were found in any of three databases: PhaSepDB, DrLLPS, or LLPSDB. Otherwise, they were classified as not in condensates. Mann-Whitney test was performed on the datasets with * meaning p < 0.05, ** p < 0.01.

Like SR and SR-like proteins, many phase-separating proteins contain RRM domains, such as TDP-43(53), FUS(54), and hnRNP proteins(55,56). Additionally, while RS repeats are abundant in nuclear speckles, not all proteins that localize to nuclear speckles possess RS repeats. We sought to understand what unique features make some RRM-containing proteins localize to condensates. To this end, we extended our bioinformatics analysis to all human RRM-containing proteins (Fig. 6B). Using the UniProt database to identify manually annotated RRM domains of proteins confirmed at the protein level, we found 206 proteins that contain in total 365 RRM domains (57). Not all RRM domains in this category have solved structures. However, because the RRM fold is well established, it can be predicted with high confidence by the software AlphaFold (58,59). We therefore used the AlphaFold Structure Database to obtain additional information about these RRM domains (Fig. 6B). We then searched for these proteins in three major phase separation databases to determine whether they were found in condensates (Fig. 6B). We found that RRM domains in phase-separating proteins have more acidic residues than those not found in condensates (Fig. 6C, compare columns 3 and 4). No significant composition differences were found for aromatic, basic, or Arg residues. Aromatic residues usually reside in protein hydrophobic cores. However, only solvent accessible aromatic residues can potentially mediate phase separation. Therefore, we used the CalcSA helper program in Xplor-NIH (51) in combination with AlphaFold structures to identify surface-exposed aromatic residues whose sidechain atoms were at least 60% solvent exposed(Fig. 6B). Among RRM domains that have not been found in condensates, a significant number possessed no aromatic residues with this level of solvent exposure, and the median value was one (Fig. 6D, column 1). RRM domains of proteins found in condensates possessed exposed aromatic residue with a median of two, with the majority possessing at least 1 (Fig. 6D, column 2). To determine whether proteins that contain RS repeats possess properties distinct from other phase-separating proteins, we then narrowed down our analysis to proteins containing 4-mer RS repeats (Fig. 6E). We found that proteins possessing RS repeats were especially likely to have surface-accessible aromatic residues on their RRM domains (Fig. 5E, columns 1 and 2). In the case of SRSF1, we found some acidic residues are close to aromatic residues (D80 to Y79 in Fig. 5C, E68 to F88 in Fig. 5D), which enables simultaneous formation of cation-pi stacking and salt bridges. Therefore, we examined residues within 6 Å of aromatic residues for phase-separating proteins (Fig. 6B). We found that RRM domains of RS-repeat containing proteins were more likely to have neighboring acidic residues, with the majority possessing 1-2 aromatic residues with neighboring acidic groups (Fig. 6E, columns 3 and 4). This trend was also present in proteins that localize to nuclear speckles (Fig. 6F, columns 3 and 4). Interestingly, in nuclear speckles, which are rich in RS repeats, this trend persisted even among proteins that did not possess their own 4-mer RS repeats (Fig. 6F, columns 3 and 5). This suggests that cooperation between aromatic and acidic residues may be an additional method for determining which proteins will localize to nuclear speckles and interact with RS repeats.

## DISCUSSION

There is currently a need to improve methods for determining which proteins phase separate and by what mechanism they do so, but solubility concerns make isolated experiments out of reach for many proteins. To solubilize phase-separating proteins, denaturants or high concentrations of salts are typically used. Denaturants are unsuitable for experiments characterizing native state proteins. High concentrations of salts are flawed as they interfere with NMR(44), SAXS(60) and circular dichroism(61). Some proteins experience salting out when ionic co-solutes are introduced (37,49,56). For example, we found that the solubility of SRSF1 is around 2 to 7 μM in 1 to 5 M of NaCl (data not shown). The protein solubilizing strategy used here is of general applicability, not just confined the examples of SRSF1 and Nob1. Our bioinformatic search revealed an abundance of RS-containing proteins. We further revealed a positive correlation between the length and the number of these repeats with the chance that the proteins containing them are found in condensates.

Characterization of intermolecular interactions that occur in the dispersed (soluble) state is an accepted method of understanding what interactions lead to phase separation (48,49,62). A previous comparison of intermolecular PRE spectra of the protein FUS in the dispersed versus the condensed (phase separated) state indicated that the transient intermolecular interactions seen between molecules in the solution state are comparable to the intermolecular contacts seen when the protein is phase separated ((37,54) as discussed in (48)). Our technique is unique in that it provides transient competition for the intermolecular interactions that lead to phase separation without abolishing these interactions entirely. We find that both RNA binding (Fig. S3B) and homotypic intermolecular interactions (Fig. 4E) can still occur in a peptide-containing buffer. However, the presence of the peptide weakens intermolecular interactions enough to allow high quality NMR spectra to be obtained. Here, we find that targeted competition for intermolecular interactions provides direct control over the critical point for phase separation, enabling experiments to be performed in the dispersed state that otherwise might not be possible. The ability to compare dispersed and condensed states for more proteins is still desirable. NMR spectra of isolated low complexity domains in the condensed state have been obtained successfully for several proteins including HNRNPA2, FUS, and a Caprin1-pFMRP complex (37,49,63-65). However, interactions between the structured domains and low complexity domains of these proteins have not yet been probed using these techniques. One bottleneck to obtaining usable condensed state samples involves obtaining high concentrations of soluble protein before inducing phase separation in the sample (37,49,63). Due to the high concentrations needed for an NMR backbone assignment and the negative effect of sample viscosity on NMR spectral quality, it is also more practical to perform backbone assignments of proteins in the dispersed states (37,49,63). We hope our method may serve as a useful tool in expanding these techniques.

Homotypic intermolecular interactions and interactions with the peptides involve residues similar to those involved in intramolecular interactions. However, intermolecular interactions seem to have a greater preference for the more negatively charged RRM1. Whereas interactions with RRM1 appear at all concentrations studied, a concentration of 25 mM peptide is needed to observe bleaching of W134 and A150 on the electropositive RRM2 domain (Fig. 4C). This difference may be due to the fact that, while intramolecular interactions involve an RS domain tethered to RRM2 that helps facilitate interaction, external RS repeats do not have a method of compensating for this charge repulsion.

This preference for RRM1 is interesting because the interactions seen on RRM2 involve the same residues that bind to RNA(66), but the interactions on RRM1 are opposite the RNA-binding interface (67). In fact, chemical shift perturbations performed in a previous study indicate that the α1 helical residues on RRM1, in particular, remain virtually unaffected when two different RNA ligands are introduced, indicating there are also no allosteric effects (67). Further, negative charges in this region are conserved across multiple SR protein family members (Fig. 6A). This suggests that RRM domains of SR proteins may have alternative sites used to mediate the protein-protein interactions that lead to phase separation. This is important because it means that the effect of phase separation on RNA binding can potentially be studied by disruption of these distal sites.

As we learn more about biomolecular condensates, it is of interest to understand what causes proteins to migrate to one condensate over another. It has been shown that the isolated SRSF2 RRM can localize to speckles on its own(68), which suggests there may be an additional molecular grammar within the structured components of these proteins that directs them towards nuclear speckles. We find that the RRM domains of proteins that have been found in nuclear speckles are more likely to possess solvent exposed aromatic residues surrounded by acidic residues than proteins found in different types of condensates (Fig. 6F, columns 3 and 4). This trait of RRM domains appears to be a localization-factor that can stand independently, as it accounts for some proteins that localize to nuclear speckles even though they do not possess their own RS repeats (Fig. 6F, columns 3 and 5). It is known that speckles rely on RS repeats as scaffolds, as truncation of SRRM2’s regions containing RS repeats (Fig. S1) disrupts speckles (21,29). It is possible that these RS repeats function in part by providing multiple interaction sites for this type of RRM.

### Ideas and Speculation

We demonstrate that electronegative α1 and α2 helices along with neighboring aromatic residues serve as interacting sites for unphosphorylated RS repetitive sequences. Further, our bioinformatics data shows that such combinations of aromatic and acidic residues are a unique characteristic of RRM domains found in nuclear speckles. This finding has implications for how phosphorylation might change interactions within the speckles. If negatively charged residues are important for maintaining protein-protein interactions, charge repulsion between a hyperphosphorylated tail and acidic residues might be one reason that SR proteins leave the speckles upon hyperphosphorylation(32). Phosphoserines of RS repeats have been proposed to form salt bridges with neighboring arginine residues(69), although any such contacts are likely short-lived and do not result in a stable secondary structure(70,71). Temporary phosphoserine-arginine contacts may be sufficient to compete with the highly transient cation-pi stacking interactions that we observe here.

Our bioinformatics analysis revealed three traits of RRMs that confer a greater likelihood of phase separation. The first, exposed aromatic residues (Fig. 6D), is consistent with an association between stacking interactions and phase separation that has been established in a variety of studies (37,55,72,73). The second, more acidic residues (Fig. 6C), partly aligns with our finding that the more electronegative RRM1 domain of SRSF1 formed more intermolecular contacts, although the binding pattern should be different for low complexity domains that are not RS repeats. The third, more arginine residues surrounding exposed aromatic groups (Fig. 6D), was found unexpectedly across multiple condensate types (Fig. S8), and it is worthy of future study. The function of these residues likely varies with condensate type. For instance, arginine residues on RNA binding domains in FUS proteins form cation-pi interactions with aromatic low complexity domains(74) and. Alternatively, it has been proposed that arginine-arginine stacking interactions may occur within nuclear speckles (68) based on recent evidence that these interactions occur through the charge balance of intervening water molecules (75). Arginine residues may also play a more general role of enabling binding to local aromatic residues by balancing the charges conferred by the increase in acidic residues and/or forming interactions with aromatic residues that maintain their surface exposure.

## MATERIAL AND METHODS

### SRSF1 expression and purification

The DNA encoding human SRSF1 was sub-cloned into pSMT3 using BamH I and Hind III. The ΔRS construct and SRSF1 the mutants SRSF1 C16S C148S N220C (AKA N220C), SRSF1 C16S C148S T248C (AKA T248C), and SRSF1 C16S C148S (AKA NoC) were prepared using mutagenesis PCR. All proteins were expressed by BL21-CodonPlus (DE3) cells in LB media or minimal media supplemented with proper isotopes for NMR experiments. Hyperphosphorylated SRSF1 was prepared by co-transformation of BL21-CodonPlus (DE3) cells using pSMT3/SRSF1 and CDC2-like kinase 1 (CLK1) cloned in pETDuet-1. Cells were cultured at 37 °C to reach an OD600 of 0.6, and 0.5 mM IPTG was added to induce protein expression. Cells were further cultured 16 hours at 22 °C. The cells were harvested by centrifugation (4000 RCF, 15 min). The cell pellet was re-suspended in 20 mM HEPES, pH 7.5, 150 mM Arg/Glu, 2 M NaCl, 25 mM imidazole, 0.2 mM TCEP supplemented with 1 mM PMSF, 1 mg/mL lysozyme, 1 tablet of Pierce protease inhibitor, and 1 mM NaVO4 for the hyperphosphorylated construct. After three freeze-thaw cycles, the sample was sonicated and centrifuged at 23,710 g for 40 min using a Beckman Coulter Avanti JXN26/JA20 centrifuge. The supernatant was loaded onto 5 mL of HisPur Nickel-NTA resin and then eluted with 60 mL of 20 mM MES pH 6.5, 300 mM imidazole, 600 mM Arg/Glu, and 0.2 mM TCEP. The eluted sample was cleaved with 2 µg/mL Ulp1 for 2 hours at 37 °C. The four unphosphorylated SRSF1 constructs (WT, N220C, T248C, NoC) were further purified by a 5-mL HiTrap Heparin column. The hyperphosphorylated SRSF1 was further purified by a 5-mL Cytiva Fast Flow Q column. The eluted samples from the ion exchange step were further purified by a HiLoad 16/60 Superdex 75 pg size exclusion column equilibrated with 800 mM Arg/Glu, pH 6.5, 1 mM TCEP, 0.02% NaN3. The protein identities were confirmed by mass spectrometry. As reported in previous study, 18-phosphates were added on the RS region of SRSF1(76). The protein purities were judged to be > 95% based on SDS-PAGE.

### Nop9 and Nob1 expression and purification

Nop9 and Nob1 expression and purification are detailed in published paper(77,78). SUMO-tagged proteins were induced by 0.4 mM IPTG and expressed at 22 °C overnight in E. coli strain BL21-CodonPlus (DE3). The LB miller medium was supplemented with 0.1 mM ZnSO4 for Nob1 expression. Cell pellets were re-suspended in 25 mM HEPES, pH 7.5, 1 M NaCl, 1 mM TCEP, 25 mM imidazole, 1 mg/mL lysozyme and lysed by sonication, followed by centrifugation. The supernatant was applied to HisPur Ni-NTA resin, washed with 200 mL of loading buffer, and eluted with 25 mM HEPES, pH 7.5, 500 mM NaCl, 1 mM TCEP, 500 mM imidazole. The SUMO tag was cleaved overnight with 2 µg/mL of Ulp1 at 4 °C. The cleaved sample was purified by a 5-mL HiTrap Heparin column (GE Healthcare), and polished using a HiLoad 16/60 Superdex 200 column (GE Healthcare) equilibrated in 25 mM HEPES, pH 7.5, 500 mM NaCl, and 1 mM TCEP. The protein purities were > 95% based on SDS-PAGE.

### NMR assignment

SRSF1 cultured in ^2^H,^13^C, ^15^N M9 media was concentrated to 370 µM in 100 mM ER4, 400 mM Arg/Glu, pH 6.4, 1 mM TCEP, 10% D2O, and 0.02% NaN3. Triple resonance assignment experiments HNCA, HNCACB, HN(CO)CA, HN(CO)CACB, HNCO, and HN(CA)CO were collected at 37 °C on a Bruker Avance III-HD 850 MHz spectrometer installed with a cryo-probe. Approximately 85% of the protein backbone region was assigned using this method. Another approximately 13% of backbone exists in the disordered state with highly degenerate sequences, which leads to heavy peak overlap. These RS and G-rich regions were grouped into clusters. Multiplicity selective in-phase coherence transfer (MUSIC) experiments were collected to further characterize the clusters and verify the assignment of the rest of the protein. MUSIC was performed on SRSF1 for the following amino acids: Ser, Arg, Thr, Asn, Ala, Tyr/His/Phe, Pro, Asn/Gln, Met, and Gly. When used in combination with analysis of the effect of paramagnetic tag placement, peak clusters were able to be assigned to locations on the disordered regions. The NMR data was processed using NMRPipe(79), and assignment was performed using NMRViewJ(80). The assignment of the well dispersed regions (85% of the protein) has been submitted to BMRB (ID: 51299). Additional data available upon request.

### Paramagnetic relaxation enhancement (PRE) measurements

RS peptide with a sequence “SRSRSRSRC” was synthesized and purified by GenScript with a purity > 98%. The cysteine residue at the C-terminus was introduced for MTSL labelling. RS peptide was mixed with MTSL in a molar concentration ratio of 1:4. The pH was adjusted to 7.0 before a 2-hour labelling at room temperature. To remove unreacted MTSL, 10 mL of ether was added to the sample, and the mixture was vortexed and spun at 4,000 rpm for 5 min. The extraction process was repeated twice. After purification, the pH of the peptide was adjusted to 6.5 using KOH, and the peptide was lyophilized. ^1^H paramagnetic relaxation enhancement (PRE) data was gathered at 37 °C on a Bruker Avance III-HD 850 MHz spectrometer installed with a cryo-probe. Titrations were performed by adding solid peptide to a ^15^N-labeled SRSF1 construct without cysteine (C16S/C148S) in 200 mM Arg/Glu, pH 6.3, 1 mM TCEP. PRE spectra were obtained for the protein in 200 mM Arg/Glu alone, 200 mM Arg/Glu with 2.5 mM peptide, 200 mM Arg/Glu with 25 mM peptide, and 200 mM Arg/Glu with 50 mM peptide. After the spectrum with 50 mM peptide was collected, MTSL was quenched using 10 mM sodium ascorbate, and a PRE experiment was run on the quenched sample. All spectra and additional PRE calculations are available upon request.

To prepare the sample for inter-molecular PRE, an SRSF1 construct with one cysteine at the C-terminal end of the RS tail (SRSF1, C16S/C148S/T248C) was obtained using mutagenesis PCR, and the mutated protein was expressed by BL21-CodonPlus (DE3) cells in LB media. The protein was exchanged into an MTSL labelling buffer of 0.8 M Arg/Glu, 100 mM NaCl, 50 mM Tris-HCl pH 7.0 using a HiPrep 26/10 desalting column. The sample was diluted to a concentration of 20 µM, and MTSL was added to a concentration of 0.4 mM. The sample was incubated in the dark at 37 °C for 12 hours, after which a desalting column was used to remove unreacted MTSL. The MTSL-labelled, NMR-inactive SRSF1 was mixed in a 1:1 ratio with a ^15^N SRSF1 construct with no cysteine (C16S/C148S) in a buffer of 100 mM ER4, 5 mM MES, pH 6.4, 400 mM Arg/Glu, and 5% D2O. The final concentration of protein was 420 µM (210 µM SRSF1 C16/148S T248C-MTSL and 210 µM ^15^N SRSF1 C16/148S).

To measure intramolecular PRE, an SRSF1 construct with one cysteine at the center of the first RS domain (SRSF1 C16S/C148S/N220C) was obtained using mutagenesis PCR and the purification method described above with the growth performed in M9 media containing ^15^N isotopes. MTSL was labelling was performed as described above. The final concentration of protein was 220 µM.

The low concentration intermolecular PRE was collected between SRSF1 C16/148S N220C-MTSL and ^15^N SRSF1 C16/148S at a total concentration of 185 µM (93 µM of each construct) as described above. MTSL-only PRE experiments were conducted by adding 220 µM MTSL to 220 µM ^15^N SRSF1 C16/C148S in the NMR buffer described above.

All PRE measurements were carried out using a pulse sequence developed by Junji Iwahara(81). Diamagnetic data were collected after adding 2 mM ascorbic acid. The NMR data was processed using NMRPipe(79) and analysed using NMRViewJ(80). PRE values and errors were estimated as described previously(81).

Residues were considered above noise level if their intensity at the second time point on the diamagnetic spectrum was greater than five times the standard deviation of the spectrum. Peaks below this noise level threshold were eliminated from analysis on all spectra. Peaks in regions of high spectral overlap were also eliminated. If at the first timepoint of collection (approximately 12 ms after the first 90° pulse), the intensity of the paramagnetic peak was less than or equal to half of the intensity of the diamagnetic peak at that timepoint, the peaks were defined as bleached. In these circumstances, the relaxation occurred too quickly to allow fitting of the exponential decay curve. Residues G52 and R154 met the definition of both bleached and noisy. They were eliminated from analysis.

### Solubility assays

Purified protein aliquots (40 μL) were incubated with 3.2 M ammonium sulfate on ice for 30 min before 10-minute centrifugation at 14,000 RCF at 4 °C. After confirming that no protein was present in the supernatant, the supernatant was discarded. The pellets were re-suspended in 20 µL of corresponding buffers and shaken at room temperature for 30 min. The re-suspensions were further centrifuged at room temperature at 14,000 RCF for 5 min. The protein concentrations in supernatants were measured using UV absorbance at 280 nm. Error bars represent technical repeats. The initial concentrations of full-length SRSF1 constructs and full-length snRNP70 were 250 μM and 85 μM, respectively. As RS-deleted SRSF1 has a higher solubility, the initial protein concentration used for this construct was 400 μM.

### Molecular Graphics

An Alphafold structure was downloaded from the Uniprot website for SRSF1 ΔRS and refined using Xplor-NIH. Restraints used for refinement included dihedral angles obtained from the assignment, RDC values obtained in a previous study(82), and chemical shift perturbations (data available upon request). PRE values were projected onto to the structure in PyMOL by reassigning B-factors and coloring on a ramp scale.

### MD simulations

An Alphafold structure was downloaded from the Uniprot website for SRSF1 ΔRS and refined using Xplor-NIH. Docking of peptides was accomplished with Xplor-NIH using a restrained rigid-body simulated annealing protocol refined against the PRE, CSP, RDC, and dihedral angle data (additional data available upon request). The structure was further refined using AMBER20 with the ff19SB forcefield. Solvation was performed with explicit TIP3P water molecules with 0.15 M NaCl used to balance the charges. The simulation temperature was set to 300 K, and the cutoff distance of nonbonded interactions was set to 10 Å. The simulation was run for 200 ns. Due to competition with neighboring arginine residues in the linker, a distance restraint was applied during the MD simulation to maintain contact between residue 88 and the peptide.

### Bioinformatics analysis

Domain annotations and sequences for human proteins were downloaded from the Uniprot website. Analysis was restricted to full-length, reviewed, human proteins for which there was evidence at the protein level. A Python script was used to search for consecutive Ser-Arg or Arg-Ser repeats of 4, 6, or 8 amino acids. Identification of proteins in condensates was based on databases PhaSepDB (PhaSepDB2.0 download), DrLLPs, and LLPSDB (natural protein download).

RRM domains were identified by further restricting our Uniprot search to proteins containing RRM domains of any manual assertion. In analysis of the correlation between condensation, RS repeats, and RRM domains, we found 206 proteins that together contained 365 RRM domains. Distinct RRM domains within the same protein were analyzed as separate entities. Because structures have not been solved for all 365 RRM domains analyzed, structures were downloaded from the Alphafold repository. For proteins in which multiple models were available, the F1 model was used. Xplor-NIH version 3.4 was used for surface area calculations. A series of python scripts was used to calculate solvent exposure of carbon side chain atoms. Aromatic residues (F,Y,W) were defined as “solvent exposed” if the side chain atoms possessed more than 60% of the highest possible degree of solvent exposure. “Neighboring residues” were defined as those within 6 Angstroms of a given aromatic residue based on the Alphafold structures. Statistics were analyzed using the Mann-Whitney test. Python scripts are available upon request and via our github link: https://github.com/taliafargason/Aromatic_neighbors.

### Imaging

An SRSF1 construct (SRSF1 C16S/C148S/N220C) was tagged with Maleimide-Alexa488, dissolved into 800 mM Arg/Glu, pH 6.5, 0.2 mM TCEP, and stored at a concentration of 28 µM. snRNP70 was tagged with Alexa488, dialyzed into 800 mM Arg/Glu, pH 8.4, and concentrated to 26 µM. Surplus Alexa488 dye was removed by a desalting column. The protein was then diluted to its final concentration in 100 mM KCl, 10 mM MES pH 6, 0.1 mM TCEP, with or without short peptides in a 96-well Cellvis glass bottom plate coated with Pluronics F127. Images are brightfield/GFP channel overlays taken on a Cytation5 imager using the software Gen5 3.10.

## DATA AVAILABILITY

The assignment of the well dispersed regions of SRSF1 has been submitted to BMRB (ID: 51299).

## SUPPLEMENTARY DATA

Supplementary Data are available online.

## ACKNOWLEDGEMENT

We want to thank UAB Central Alabama High-Field NMR Facility. We also want to acknowledge Dr. Jinfa Ying in Ad Bax lab in at NIDDK, Dr. Charles D. Schwieters at NIH for technical support. This work is supported by U.S. National Science Foundation, MCB 2024964.

## FUNDING

This work was supported by the U.S National Science Foundation [MCB2024964 to J.Z.]. Funding for open access charge: National Science Foundation.

## CONFLICT OF INTEREST

None declared

## Notes

https://github.com/taliafargason/Aromatic_neighbors

### Competing Interest Statement

The authors have declared no competing interest.

### Summary of Updates

Figure added to the supplemental.

