## Supplemental Materials for "Peptides that Mimic RS repeats modulate phase separation of SRSF1, revealing a reliance on combined stacking and electrostatic interactions"

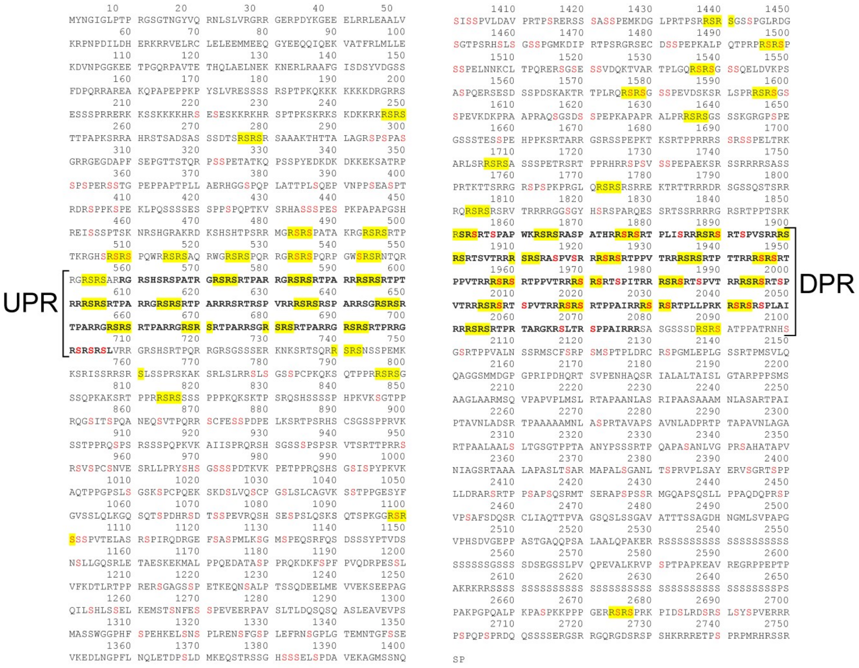


Figure S1: Amino acid sequence of the protein SRRM2 with 56 4-mer RS repeats highlighted in yellow. Serines in red can be phosphorylated according to the Uniprot database. Deletion of the undeca-repeat (UPR) and dodeca-repeat (DPR) regions (indicated by bold font and brackets) results in dissociation of nuclear speckles(1).

A

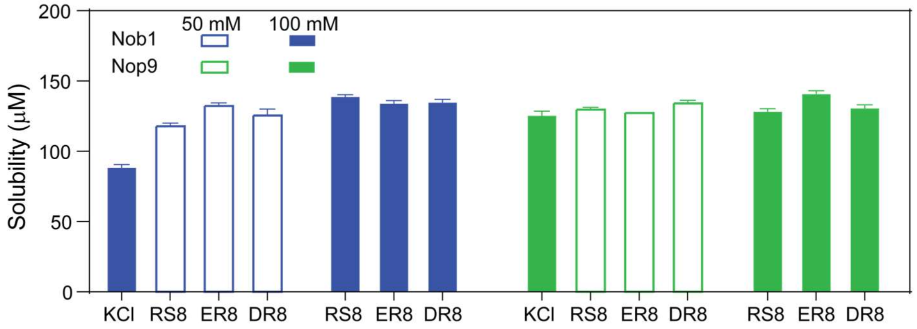


B

Nob1 sequence:

10 20 30 40 50
**MTENQTAHVR** ALILDATPLI TQSYTHYQNY AQSFYTTPTV FQEIKDAQAR
60 70 80 90 100
KNLEIWQSLG TLKLVHPSEN SIAKVSTFAK LTGDYSVLSA NDLHILALTY
110 120 130 140 150
ELEIKLNNGD WRLRK**KPGDA LDASKADVGT DGKQKLTEDN KKEEDSESVP**
160 170 180 190 200
**KKKNKRRGGK KQKAKREARE AREAENANLE LESKAEEHVE EAGSKEQICN**
210 220 230 240 250
**DENIKESSDL NEVFEDADDD GDWITPENLT EAIIKDSGED TTGSLGVEAS**
260 270 280 290 300
**EEDRHVALNR PENQVALATG DFAVQNVALQ MNLNLMNFMS GLKIKRIRNY**
310 320 330 340 350
**MLRCHACFKI FPLPKDGKPK HFCASCGGQG TLLRCAVSVD SRTGNVTPHL**
360 370 380 390 400
**KSNFQWNNRG NRYSVASPLS KNSQKRYGKK GHVHSKPQEN VILREDQKEY**
410 420 430 440 450
**EKVIKQEEWT RRHNEKILNN WIGGGSADNY ISPFAITGLK QHNVRIGKGR**

**YVNSSKRRS**

C

Nop9 sequence:

10 20 30 40 50
**MGKTKTRGRR HQDKQRKDEF EPSSNSAKEH IQQEESTYND EAE**IKETQPQ
60 70 80 90 100
MFFGVLDREE LEYFKQAEST LQLDAFEAPE EKFQFVTSII EEAKGKELKL
110 120 130 140 150
VTSQITSKLM ERVILECDET QLKDIFQSFN GVFFGLSCHK YASHVLETLF
160 170 180 190 200
VRSAALVERE LLTPSFDNNE KEGPYVTMEN MFLFMLNELK PHLKTMMNHQ
210 220 230 240 250
YASHVLRLLI LILSSKTLPN STKANSTLRS KKSKIARKMI DIKDNDDFNK
260 270 280 290 300
VYQTPESFKS ELRDIITTLY KGFTNGAESR SDISQSTITK FREYSVDKVA
310 320 330 340 350
SPVIQLIIQV EGIFDRDRSF WRLVFNTADE KDPKEESFLE YLLSDPVGSH
360 370 380 390 400
FLENVIGSAR LKYVERLYRL YMKDRIVKLA KRDTTGAFVV RALLEHLKEK
410 420 430 440 450
DVKQILDAVV PELSMLLNSN MDFGTAIINA SNKQGGYLRD DVIAQLIQKY
460 470 480 490 500
YPEKSDAKNI LESCLLLSAS TLGNTRDDWP TAEERRRSVF LEQLIDYDDK
510 520 530 540 550
FLNITIDSML ALPEERLIQM CYHGVFSHVV EHVLQTTRVD IIKRKMLLNI
560 570 580 590 600
LSKESVNLAC NVYGSHIMDK LWEFTAKLTL YKERIARALV LETEKVKNSI
610 620 630 640 650
YGRQVWKNWK LELYVRKMWD WKKLIKEQEF EIFPNSKPLQ PKPEKH**SRER**
660
**NNSKEGSAFK KQKHYR**

Figure S2: Repetitive peptides solubilize other proteins containing similar repetitive regions (A) Solubilizing effects of 8-mer Arg-Ser (RS8), Glu-Arg (ER8), and Asp-Arg (DR8) were tested on Nob1 (blue bars) and Nop9 (green bars) at peptide concentrations of 50 mM (open bars) and 100 mM (filled bars). A buffer of 100 mM KCl was used as a control. (B) Nob1 protein sequence. (C) Nop9 protein sequence. Unstructured regions are shown in bold fonts. The basic and acidic-basic residue regions are in cyan and red, respectively.

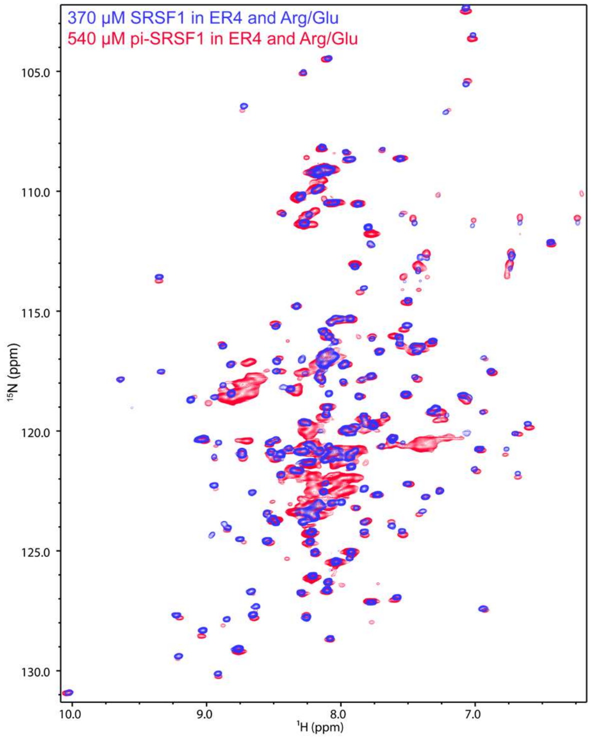


Figure S3: The buffer 100 mM ER4, 400 mM Arg/Glu, pH 6.4 was effective at solubilizing and optimizing spectral quality for both unphosphorylated and hyperphosphorylated SRSF1 at high enough concentrations for NMR assignment. $^{15}$N-TROSY-HSQC overlay of 370 µM SRSF1 (blue) and 540 µM phosphorylated SRSF1 (pi-SRSF1, red) in this buffer.

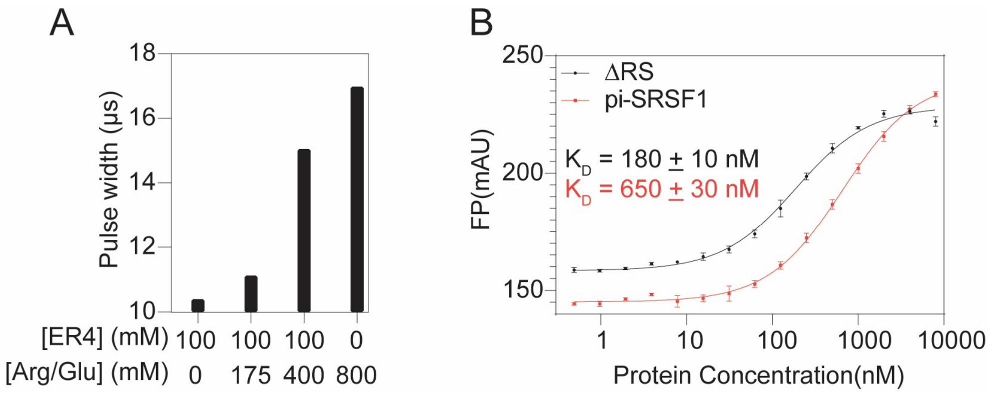


Figure S4: The buffer 100 mM ER4, 400 mM Arg/Glu, pH 6.4 provided a usable pulse width and did not abolish ligand binding. A) Pulse width of various concentrations of ER4 and Arg/Glu. B) RNA-binding was weakened but not abolished by the presence of peptides in the binding buffer. Fluorescence polarization assays for binding of SRSF1 constructs with an Alexa488-tagged, 10 nM RNA probe (UCAGAGGA). Both binding assays were performed in 100 mM ER4, 400 mM Arg/Glu, pH 6.5. Error bars represent SEM.

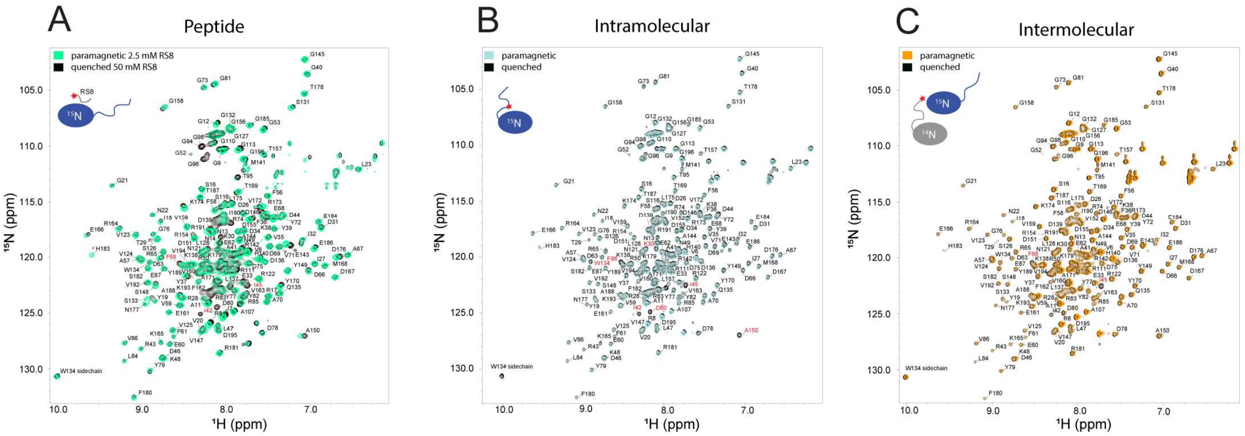


Figure S5: $^{15}$N-TROSY-HSQC overlay of samples used in A) Peptide-SRSF1, B) Intramolecular, and C) Intermolecular PRE experiments. Spectra in which the paramagnetic center was active are color coded, and spectra in which the paramagnetic center was quenched with ascorbic acid are in black. Residues marked in red came in close enough contact to paramagnetic centers that PRE could not be quantified. We refer to these residues as bleached.

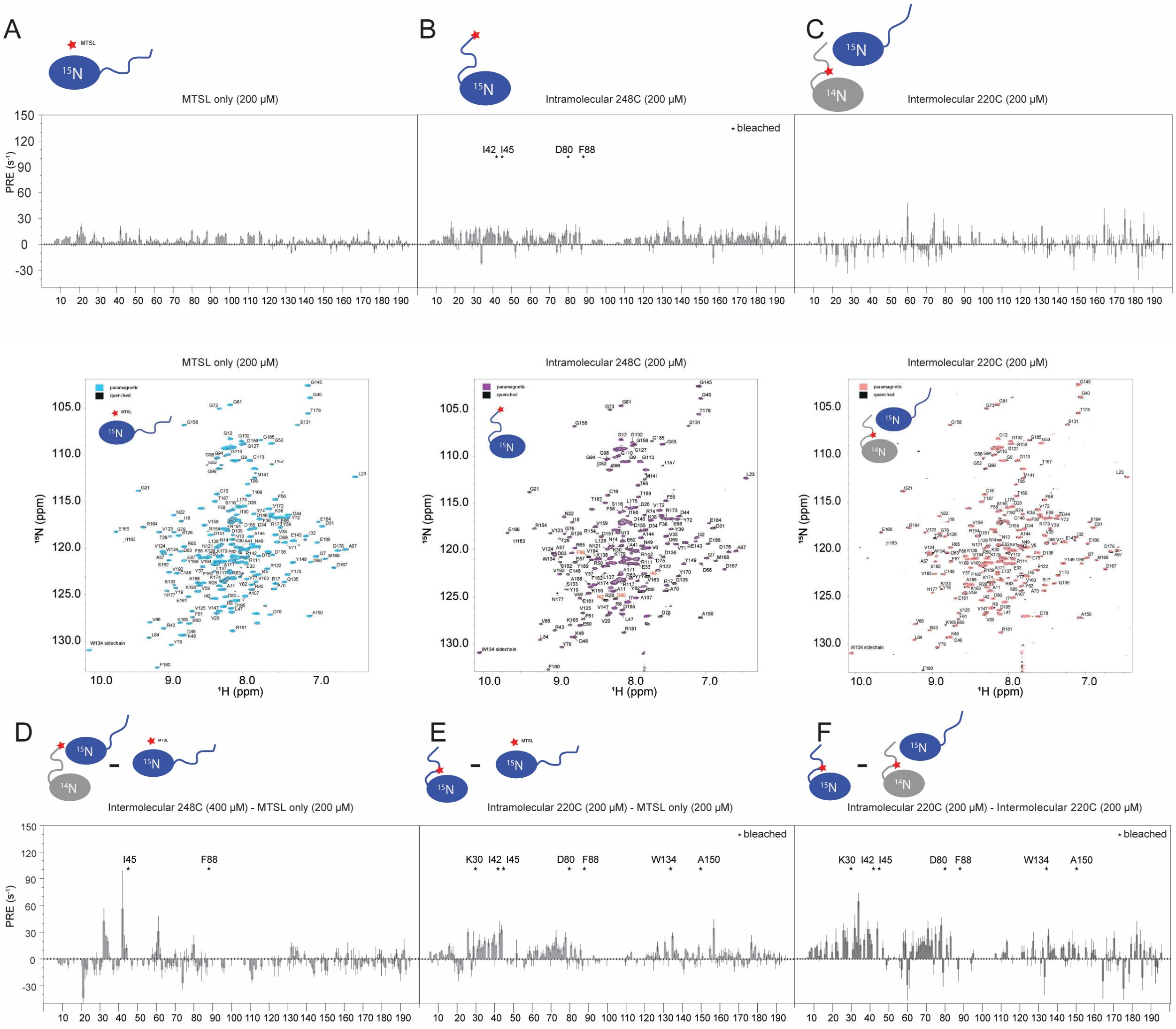

Figure S6: PRE of MTSL alone (A), an intramolecular PRE experiment with the MTSL tag on the very C-terminal end of the protein (B), and an intermolecular PRE experiment replicating the conditions of the intermolecular interactions experiment (C). When these spectra are subtracted from the experiments displayed in the main text(D-F) the trend is still similar.

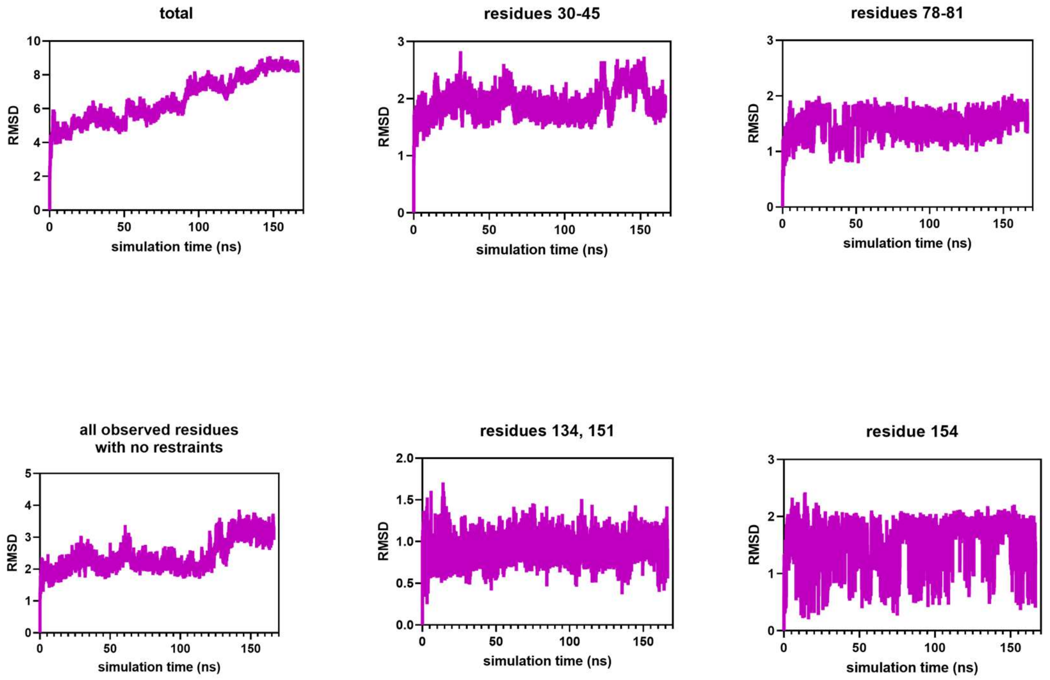


Figure S7: MD simulation trajectories of SRSF1 ΔRS interacting with RS8 peptides in a simulation in which no restraints are applied. Root mean standard deviation between the simulation starting structure and the structure at a given simulation time indicates that, in most residues under observation, an equilibrium is reached between 120-180 ns that was used for observation. Numbers above the individual plots indicate the residues with which a given peptide interacts.

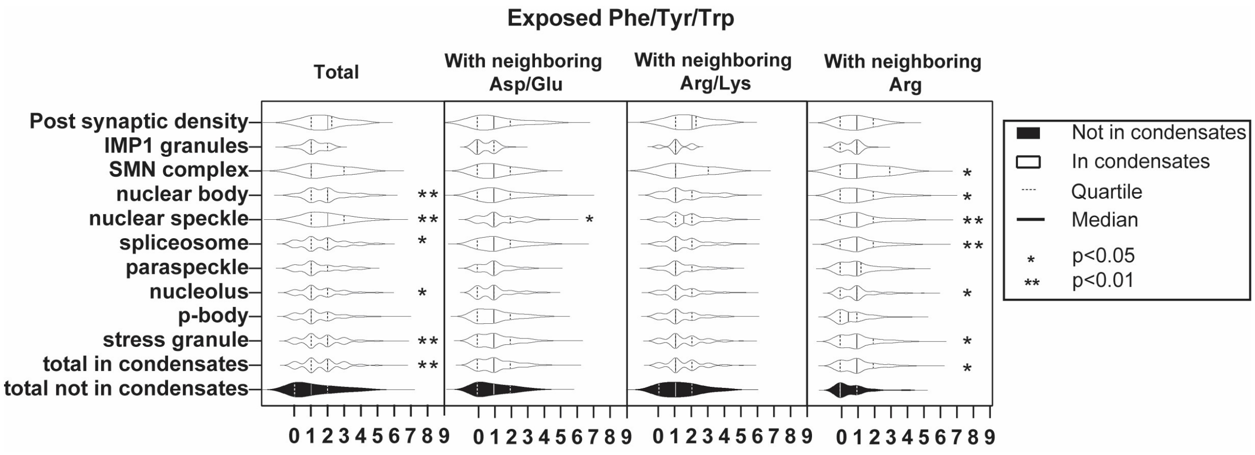


Figure S8: Characterization of surface exposed aromatic residues on RRM domains based on condensate type. p-values were calculated using the Mann-Whitney test.

1. Xu, S., Lai, S.K., Sim, D.Y., Ang, W.S.L., Li, H.Y. and Roca, X. (2022) SRRM2 organizes splicing condensates to regulate alternative splicing. *Nucleic Acids Res*, **50**, 8599-8614.
